# Supervised Machine Learning for Bioelectrical Cellular Networks

**DOI:** 10.1101/2024.04.30.591880

**Authors:** Rajeev Jaundoo, Travis J.A. Craddock, Jack A. Tuszynski

**Affiliations:** University of Alberta; Nova Southeastern University

## Abstract

Cells utilize bioelectricity to form networks as well as regulate and control a variety of processes such as apoptosis, tumor suppression, and voltage-gated ion channels. In-silico modeling of bioelectrical networks can be performed using BETSE, an application that models gap junctions and ion channel activity of networked cells, but its usage of matrix-based differential equations to estimate these properties limits simulations based on the amount of computational resources available. To alleviate this issue, we trained a total of 8 machine learning models to replace three core functions of BETSE, that is, 1) predicting the average transmembrane potential (V_mem_) of an entire cellular network, 2) predicting the V_mem_ of each individual cell within the network, and finally, 3) predicting the average ion concentrations of sodium, potassium, chloride, and calcium within the cell network. For objective 1, the random forest model was shown to be most performant over all 4 scoring metrics, in objective 2 both the decision tree and k-nearest neighbors models scored best in half of all metrics, and for objective 3 the super learner, a meta-learner comprised of multiple base learners, scored best among all scoring metrics. Overall, these models provide a more resource efficient method of predicting properties of bioelectric cellular networks, and future work will include further properties such as temperature and pressure.

## 2. Introduction

Bioelectricity is used by cells to communicate with one another and regulate various processes including voltage-gated ion channels, tumor suppression, and apoptosis among others (Pietak & Levin, 2016; Srivastava et al., 2021). Bioelectrical cellular networks can be modeled in-silico using the Bio Electric Tissue Simulation Engine (BETSE), an application that simulates gap junctions (GJs) and ion channel activity of non-neuronal cells using matrix-based differential equations. This allow BETSE to estimate the cytosolic and extra-cellular concentrations of sodium (Na^+^), potassium (K^+^), chloride (Cl^-^), and calcium (Ca^2+^) ions, as well as each cell’s transmembrane potential (V_mem_), or the difference in electrical potential between the intra-cellular space and extra-cellular medium (Pietak & Levin, 2016; Pietak & Levin, 2017). Moreover, BETSE has been validated to predict both the V_mem_ as well as intra-cellular ion concentrations of Xenopus oocytes with only <10% difference from experimental values (Pietak & Levin, 2016).

The V_mem_ of a cell, measured in millivolts (mV), affects, and in turn is affected by, ion concentration. For example, if a cell contains more Cl^-^ (negative) than Na^+^ (positive), then its V_mem_ will be more polarized, and vice versa. Interestingly, V_mem_ can also be used as a marker for cancer because tumor cells found in various parts of the body including the immune and reproductive systems, lungs, thyroid, and stomach/intestine, are typically more depolarized compared to their healthy counterparts (Srivastava et al., 2021; Tuszynski et al. 2017). Not only that, but Tuszynski et al. (2017) have shown in-vitro that artificial polarization of salamander tumor cells back to a healthy V_mem_ can prevent and even reverse tumorigenesis in some cases. The same was true for the reverse, where depolarization of healthy frog pigment cells to a cancer state V_mem_ led to drastic changes in their shape and behavior, where they became highly migratory and invaded blood vessels for instance (Tuszynski et al., 2017).

### 2.1. Objectives

The use of matrix-based differential equation solvers limits BETSE’s ability to model a bioelectrical cellular network based on available CPU power and memory. V_mem_ is calculated using the Maxwell Capacitance matrix, the Nernst-Planck equation is used to estimate ion concentration within GJs, and ion pumps are modeled using Michaelis-Menton enzyme kinetic relations (Pietak & Levin, 2016). The use of trained machine learning (ML) models would help mitigate computational limitations and allow larger networks to be predicted. In this study, we developed a set of supervised ML models to replace the core functionality of BETSE, that is:

1. Predict the average V_mem_ of an entire cellular network at a given time.
2. Predict the Vmem of each cell within a network at a given time.
3. Predict the average Na^+^, K^+^, Cl^-^, and Ca^2+^ concentrations of a network at a given time.

Where the time refers to the number of seconds of in BETSE simulation, e.g., predicting the V_mem_ of the network at 5 seconds.

## 3. Methods

First, BETSE simulations were first run to generate data for training and validation. Next, the scikit-learn Python library (Pedregosa et al., 2011) was utilized to train the following 7 learners: 1. Bayesian ridge (BAYRID), 2. decision tree (DECTRE), 3. k-nearest neighbors (KNN), 4. linear regression (LINREG), 5. feed-forward neural network (NN), 6. random forest (RANFOR), and 7. support vector regression (SVR). Additionally, the mlens Python library (Flennerhag, 2018) was used to train a super learner (SUPLRN) model, which a meta-learner comprised of various base learners, in this case we selected DECTRE, LINREG, NN, RANFOR, and SVR; see van der Laan et al. (2007). The final performance of all models was assessed using the validation set and the following 4 scoring metrics: 1. L1 loss/mean absolute error (MAE), 2. L2 loss/mean squared error (MSE), 3. median absolute error (MEDAE), and 4. Pearson R (R^2^).

### 3.1. BETSE

A YAML formatted file is used to set all parameters for a given BETSE simulation, including total initialization and simulation time (measured in seconds), starting intra-cellular and extra-cellular concentrations of Na^+^, K^+^, Cl^-^, and Ca^2+^, environment grid size, and so on. Python scripts were written to generate BETSE configuration files with randomized values for these parameters, which provided data from various conditions to train and validate the ML models on; see Table 1. The output of a BETSE simulation provides the average V_mem_ of the network and each cell at every timestep (e.g., 1 second, 2 seconds, etc.), as well as the average concentration of Na^+^, K^+^, Cl^-^, and Ca^2+^ at each timestep.

**Table 1:** Parameters that were randomized within each BETSE configuration file.

| *Option* | *Description* |
| --- | --- |
| comp grid size | Size of the environment grid |
| cytosolic Na<sup>+</sup> concentration | Intracellular sodium ion concentration |
| cytosolic K<sup>+</sup> concentration | Intracellular potassium ion concentration |
| cytosolic Cl<sup>-</sup> concentration | Intracellular chloride ion concentration |
| cytosolic Ca<sup>2+</sup> concentration | Intracellular calcium ion concentration |
| Dm_Na | Transmembrane diffusion constant for the sodium ion |
| Dm_K | Transmembrane diffusion constant for the potassium ion |
| Dm_Cl | Transmembrane diffusion constant for the chloride ion |
| Dm_Ca | Transmembrane diffusion constant for the calcium ion |
| extracellular Na<sup>+</sup> concentration | Extracellular sodium ion concentration |
| extracellular K<sup>+</sup> concentration | Extracellular potassium ion concentration |
| extracellular Cl<sup>-</sup> concentration | Extracellular chloride ion concentration |
| extracellular Ca<sup>2+</sup> concentration | Extracellular calcium ion concentration |
| time step | Time step for the initialization phase |
| time step | Time step for the simulation phase |

Note that a given BETSE simulation first involved an initialization phase that generated the cell cluster and brought the network to cell resting V_mem_. Additionally, all simulations ran for a total of 100 seconds to ensure the same amount of data would be available for every randomized condition.

### 3.2. Machine Learning

The training and validation datasets for objective 1 stemmed from a total of 14,891 BETSE simulations, while the training and validation sets for objectives 2 and 3 were based on 26,924 BETSE simulations. In all objectives 50% of the total BETSE data was used for training while the other half was set aside for validation. Table 2, Table 3, and Table 4 contain the features used to train all learners for each objective, where a description of the feature is provided along with its source, or where the feature originated from.

**Table 2:** Features used to train models for objective 1, V_mem_ prediction of the entire cell network.

| *Features* | *Feature Source* |
| --- | --- |
| Time | BETSE |
| Environmental Grid Size | Configuration File |
| Intra-cellular $\text{Na}^+$ Concentration | Configuration File |
| Intra-cellular $\text{K}^+$ Concentration | Configuration File |
| Intra-cellular $\text{Cl}^-$ Concentration | Configuration File |
| Intra-cellular $\text{Ca}^{2+}$ Concentration | Configuration File |
| Extra-cellular $\text{Na}^+$ Concentration | Configuration File |
| Extra-cellular $\text{K}^+$ Concentration | Configuration File |
| Extra-cellular $\text{Cl}^-$ Concentration | Configuration File |
| Extra-cellular $\text{Ca}^{2+}$ Concentration | Configuration File |
| $\text{Na}^+$ Diffusion Constant | Configuration File |
| $\text{K}^+$ Diffusion Constant | Configuration File |
| $\text{Cl}^-$ Diffusion Constant | Configuration File |
| $\text{Ca}^{2+}$ Diffusion Constant | Configuration File |
| *Target* | *Target Source* |
| $V_{\text{mem}}$ | BETSE |

**Table 3:** Features used to train models for objective 2, V_mem_ prediction of each cell within the network.

| *Features* | *Feature Source* |
| --- | --- |
| Time | BETSE |
| Numerical Identifier of Current Cell | BETSE |
| X-Coordinates of Cell | BETSE |
| Y-Coordinates of Cell | BETSE |
| Environmental Grid Size | Configuration File |
| Intra-cellular $\text{Na}^+$ Concentration | Configuration File |
| Intra-cellular $\text{K}^+$ Concentration | Configuration File |
| Intra-cellular $\text{Cl}^-$ Concentration | Configuration File |
| Intra-cellular $\text{Ca}^{2+}$ Concentration | Configuration File |
| Extra-cellular $\text{Na}^+$ Concentration | Configuration File |
| Extra-cellular $\text{K}^+$ Concentration | Configuration File |
| Extra-cellular $\text{Cl}^-$ Concentration | Configuration File |
| Extra-cellular $\text{Ca}^{2+}$ Concentration | Configuration File |
| $\text{Na}^+$ Diffusion Constant | Configuration File |
| $\text{K}^+$ Diffusion Constant | Configuration File |
| $\text{Cl}^-$ Diffusion Constant | Configuration File |
| $\text{Ca}^{2+}$ Diffusion Constant | Configuration File |
| *Target* | *Target Source* |
| $V_{\text{mem}}$ | BETSE |

**Table 4:** Features used to train models for objective 3, ion channel concentration prediction of the entire cell network.

| *Features* | *Feature Source* |
| --- | --- |
| Time | BETSE |
| $V_{\text{mem}}$ | BETSE |
| Environmental Grid Size | Configuration File |
| Intra-cellular $\text{Na}^+$ Concentration | Configuration File |
| Intra-cellular $\text{K}^+$ Concentration | Configuration File |
| Intra-cellular $\text{Cl}^-$ Concentration | Configuration File |
| Intra-cellular $\text{Ca}^{2+}$ Concentration | Configuration File |
| Extra-cellular $\text{Na}^+$ Concentration | Configuration File |
| Extra-cellular $\text{K}^+$ Concentration | Configuration File |
| Extra-cellular $\text{Cl}^-$ Concentration | Configuration File |
| Extra-cellular $\text{Ca}^{2+}$ Concentration | Configuration File |
| $\text{Na}^+$ Diffusion Constant | Configuration File |
| $\text{K}^+$ Diffusion Constant | Configuration File |
| $\text{Cl}^-$ Diffusion Constant | Configuration File |
| $\text{Ca}^{2+}$ Diffusion Constant | Configuration File |
| *Target* | *Target Source* |
| Intra-Cellular $\text{Na}^+$ Concentration | BETSE |
| Intra-Cellular $\text{K}^+$ Concentration | BETSE |
| Intra-Cellular $\text{Cl}^-$ Concentration | BETSE |
| Intra-Cellular $\text{Ca}^{2+}$ Concentration | BETSE |

Time is a feature that corresponds to the V_mem_ or intra-/extra-cellular concentration of Na^+^, K^+^, Cl^-^, or Ca^2+^ at a given timepoint within the BETSE simulation. For example, at 1 second the average V_mem_ of the entire cellular network may be -4.32 mV, or at 30 seconds the average extra-cellular K^+^ is 95.11 mmol/L. Objective 2 included data from individual cells at each second of the BETSE simulation, so the identifier of each cell along with their X-coordinates and Y-coordinates were included as features. For instance, at 3 seconds in the simulation, cell 1 at location (68.03, 17.41) had a V_mem_ of -4.10 mV, cell 2 at (85.29, 16.77) had a V_mem_ of -4.07, etc.

Objective 3 involved predicting multiple outputs, that is, the average ion concentrations of Na^+^, K^+^, Cl^-^, and Ca^2+^, so the RegressorChain module from scikit-learn was used to accomplish this task. Here, the first target (e.g., Na^+^) is predicted based on features within the training data, same as in objectives 1 and 2, while all subsequent targets (e.g., K^+^, Cl^-^, Ca^2+^) were predicted based on the values of all previous targets. In other words, the predicted value of Na^+^ was first predicted using only features within the training data, then the value of K^+^ was based on both features and the predicted value of Na^+^, Cl^-^ was predicted from features, Na^+^, and K^+^, and so on. Note that the order of this regressor chain was determined from the columns of the training data, from first to last: Na^+^, K^+^, Cl^-^, and Ca^2+^.

### 3.3. Validation

The performance of all models was assessed using the corresponding validation dataset for each objective, where the MAE, MSE, MEDAE, and R^2^ metrics were employed. Each of these metrics had their own advantages and caveats, for example, the use of the squared error in MSE amplified outliers, which was a potential disadvantage none of the other metrics shared. In comparison, the use of absolute error in MAE and median error in MEDAE made them more robust when dealing with outlier values. R^2^ on the other hand is a measure of how well the independent variables, the features, explain the amount of variance within the model. Although the range of this metric is (-∞, 1], anything ≤ 0 signified a model was unable to explain any of the variance between the independent and dependent variables (Chicco et al., 2021). Consequently, the range of R^2^ was considered [0,1], where all negative values were simplified to 0 without loss of meaning (Chicco et al., 2021).

## 4. Results

The performance of all models for each objective are shown in Table 5, Table 6, and Table 7. Furthermore, the R^2^ performance of all learners within the SUPLRN for objectives 1, 2, and 3 are shown in Table 8.

**Table 5:** Objective 1 performance of all models over all metrics. Highlighted entries signify the model with the best performance for that metric/column.

| *Model* | *MAE* | *MSE* | *MEDAE* | *R<sup>2</sup>* |
| --- | --- | --- | --- | --- |
| BAYRID | 21.91 | 936.46 | 16.60 | 0.57 |
| DECTRE | 2.35 | 67.00 | 0.00 | 0.97 |
| KNN | 2.25 | 56.10 | 0.00 | 0.97 |
| LINREG | 21.91 | 936.46 | 16.60 | 0.57 |
| NN | 4.24 | 64.72 | 1.40 | 0.97 |
| RANFOR | 2.07 | 48.84 | 0.00 | 0.98 |
| SVR | 6.49 | 123.32 | 3.44 | 0.94 |
| SUPLRN | 3.15 | 53.81 | 0.76 | 0.98 |

**Table 6:**
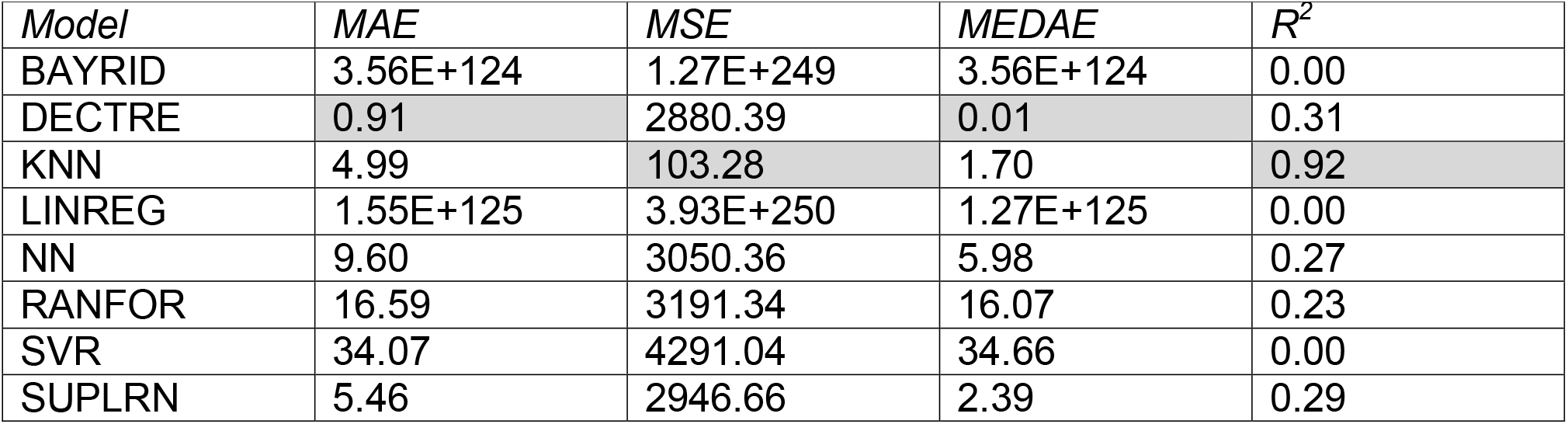
Objective 2 performance of all models over all metrics. Highlighted entries signify the model with the best performance for that metric/column. The following parameters were modified from their defaults: NN’s max_iter was set to 99,999 from 200; SVR’s max_iter was set to 75,000 from -1 (no limit); Within the SUPLRN: DECTRE’s max_depth was set to 50 from None (no limit); RANFOR’s max_depth was set to 50 from None (no limit); SVR’s max_iter was set to 50,000; NN’s max_iter was set to 3,000.

| *Model* | *MAE* | *MSE* | *MEDAE* | *R<sup>2</sup>* |
| --- | --- | --- | --- | --- |
| BAYRID | 3.56E+124 | 1.27E+249 | 3.56E+124 | 0.00 |
| DECTRE | 0.91 | 2880.39 | 0.01 | 0.31 |
| KNN | 4.99 | 103.28 | 1.70 | 0.92 |
| LINREG | 1.55E+125 | 3.93E+250 | 1.27E+125 | 0.00 |
| NN | 9.60 | 3050.36 | 5.98 | 0.27 |
| RANFOR | 16.59 | 3191.34 | 16.07 | 0.23 |
| SVR | 34.07 | 4291.04 | 34.66 | 0.00 |
| SUPLRN | 5.46 | 2946.66 | 2.39 | 0.29 |

**Table 7:** Objective 3 performance of all models over all metrics. Highlighted entries signify the model with the best performance for that metric/column. The following parameters were modified from their defaults: NN’s max_iter was set to 99,999 from 200; SVR’s max_iter was set to 250,000 from -1 (no limit). Within the SUPLRN: NN’s max_iter was set to 3,000; RANFOR’s max_depth was set to 50 from None (no limit); SVR’s max_iter was set to 250,000.

| *Model* | *MAE* | *MSE* | *MEDAE* | *R2* |
| --- | --- | --- | --- | --- |
| BAYRID | 2.88 | 19.45 | 1.98 | 0.97 |
| DECTRE | 0.22 | 0.90 | 0.06 | 1.00 |
| KNN | 0.65 | 3.33 | 0.06 | 1.00 |
| LINREG | 2.88 | 19.45 | 1.98 | 0.97 |
| NN | 1.57 | 6.58 | 0.91 | 0.99 |
| RANFOR | 0.20 | 0.57 | 0.03 | 1.00 |
| SVR | 3.25 | 19.64 | 2.80 | 0.97 |
| SUPLRN | 0.17 | 0.47 | 0.04 | 1.00 |

**Table 8:** R^2^ performance of each learner within the SUPLRN on 10-fold CV for every objective.

| *Learner* | *Objective 1* | *Objective 2* | *Objective 3* |
| --- | --- | --- | --- |
| DECTRE | 0.97 | 0.86 | 1.00 |
| LINREG | 0.57 | 0.38 | 0.97 |
| NN | 0.96 | 0.87 | 0.99 |
| RANFOR | 0.98 | 0.92 | 1.00 |
| SVR | 0.94 | 0.30 | 0.98 |

## 5. Discussion

A total of 8 different models were generated for each objective and their performance across multiple scoring metrics was compared. Objective 1 was to predict the average V_mem_ of an entire cellular network, objective 2 was to predict the V_mem_ of each individual cell within a network, and finally, objective 3 was to predict the average ion concentrations of Na^+^, K^+^, Cl^-^, and Ca^2+^ within a given cell network. A diverse set of supervised regression learners were trained, including tree-based learners such as DECTRE and RANFOR, the linear BAYRID and LINREG learners, neural network (i.e., feed-forward NN), support vector (i.e., SVR), nearest neighbor (i.e., KNN), and the SUPLRN, a meta-learner comprised of DECTRE, LINREG, NN, RANFOR, and SVR. All trained models for each objective were compared using a 2-tailed paired samples t-test, where p < 0.05 demonstrated the predicted values of 2 models were significantly different.

In objective 1, RANFOR was shown to have the best performance over all metrics, although DECTRE and KNN had the same MEDAE score and SUPLRN shared the same R^2^ value; see Table 5. Conversely, both BAYRID and LINREG performed poorly compared to the other models. While the performance of the RANFOR and KNN models (*p* = 0.00) as well as RANFOR and SUPLRN (p= 0.00) were found to be significantly different to one another, RANFOR and DECTRE (*p* = 0.30) were not. Furthermore, every pair of the DECTRE, KNN, and SUPLRN models (*p* = 0.00) were shown to have significant differences in their performances. Outside of the linear BAYRID and LINREG models, SVR also had a higher MSE score than the others, which coupled with its higher MAE and MEDAE scores, indicated all its predicted values tended to be further away from ground truth compared to DECTRE or KNN for instance. The performance of all learners within the SUPLRN are shown in Table 8, column 1. Here, the performance of each base learner reflected its performance as an individual model, where RANFOR and DECTRE had the best R^2^ scores, 0.98 and 0.97 respectively, and LINREG with the worst, 0.57.

For objective 2, Table 6 shows DECTRE had the best MAE and MEDAE scores while KNN was most performant according to the MSE and R^2^ metrics. A t-test between these models found no significant difference between their predicted outputs (*p* = 0.36). The BAYRID and LINREG models had the worst performance again, demonstrating that V_mem_ prediction was not a linear task. There was also a greater difference between the DECTRE and RANFOR models, the performance issues of the latter most likely due to the maximum tree depth hyperparameter that was capped to 50 from its default of unlimited. This was required due to computational constraints of memory. In regard to the SUPLRN, Table 8, column 2 shows RANFOR had the best R^2^ score at 0.92, followed by NN and DECTRE, 0.87 and 0.86 respectively, then LINREG at 0.38, and the worst base learner was SVR with a score of 0.30.

Finally, in objective 3 the SUPLRN model had the best MAE and MSE scores while RANFOR marginally had the best MEDAE score, 0.03 versus 0.04 for SUPLRN. The R^2^ scores were equivalent for the DECTRE, RANFOR, and SUPLRN models. Table 7 shows that SVR was the worst performing model in this objective, where it had the highest/worst MAE, MSE, and MEDAE scores and tied with BAYRID and LINREG for the lowest/worst R^2^ value. T-tests between SUPLRN and the DECTRE, KNN, and RANFOR models were *p* = 0.00, which demonstrated there was a significant difference between the performance of SUPLRN and each of these models. Similarly, DECTRE and KNN (*p* = 0.01) as well as RANFOR and KNN (*p* = 0.04) were also found to be statistically significant, while the predicted values between DECTRE and RANFOR were not (*p* = 0.11). Within the SUPLRN, LINREG and SVR had the worst R^2^ scores at 0.97 and 0.98 respectively, while DECTRE and RANFOR tied for best with a perfect rounded R^2^ score of 1.00; see Table 8, column 3.

## 6. Conclusion

The goal of this study was to develop ML models that replaced the core functionality of BETSE, an application that models bioelectrical cell networks via GJs and the ion channel activity of Na^+^, K^+^, Cl^-^, and Ca^2^ within a network. Bioelectricity plays a critical role in biology; for example, V_mem_ is used by cells to communicate with one another as well as regulate and control processes including tumor growth, stem cell differentiation, and voltage-gated ion channels among others (Pietak & Levin, 2016; Silver & Nelson, 2018). Additionally, the ability to configure BETSE parameters such as initial intra/extra-cellular Na^+^, K^+^, Cl^-^, Ca^2+^ concentrations as well as their membrane diffusion constants is useful in modeling disease cell networks such as cancer, which is known to up and down-regulate ion channels in order to proliferate (Haworth & Brackenbury, 2019).

All models were trained and validated using data generated from BETSE, where certain parameters including the size of the environmental grid and time step of the initialization and simulation phases were randomized. The use of in-silico data is the biggest limitation of this study, however, at the time of writing there was a sparse amount of experimental bioelectric data available. Furthermore, the implementation of some learners in scikit-learn was another limitation because many of them required the entire training/validation set to be loaded into memory at once. This led to issues concerning computational resources, so in objectives 2 and 3 the hyperparameters of NN, RANFOR, SVR, etc. had to be modified for training to be completed successfully.

In conclusion, for average V_mem_ prediction of an entire cellular network, the RANFOR model performed best over metrics, while V_mem_ prediction of individual cells was best performed by the DECTRE and KNN models, where each was most performant in half of all scoring metrics (and were also shown to not be statistically significant to one another either), and to predict ion concentrations, the SUPLRN model had the lowest/best MAE and MSE scores along with an equivalent R^2^ score to DECTRE, KNN, and RANFOR. Future work will include other simulation parameters such as temperature, pressure, and surface area of GJs for instance. Overall, the ML models generated in this study allow the properties of various bioelectrical cellular networks to be predicted while only requiring a fraction of the computational power needed to complete a single BETSE simulation.

